# Beat the heat: *Culex quinquefasciatus* regulates its body temperature during blood-feeding

**DOI:** 10.1101/2020.07.07.190157

**Authors:** Joanna M. Reinhold, Ryan Shaw, Chloé Lahondère

**Author notes:** To whom all correspondence should be addressed: Chloé Lahondère, Department of Biochemistry, Virginia Polytechnic Institute and State University, Blacksburg, VA, 24061, USA.

## Abstract

Mosquitoes are regarded as one of the most dangerous animals on earth. As they are responsible for the spread of a wide range of both human and animal diseases, research of the underlying mechanisms of their feeding behavior and physiology is critical. Among disease vector mosquitoes, *Culex quinquefasciatus*, which is a known carrier of West Nile virus and Western Equine Encephalitis, remains relatively understudied. As blood sucking insects, adaptations (either at the molecular or physiological level) while feeding on warm blood is crucial to their survival, as overheating can result in death due to heat stress. Our research aims to study how *Cx. quinquefasciatus* copes with heat associated with the ingestion of a warm blood-meal and to possibly uncover the adaptations this species uses to avoid thermal stress. Through the use of thermographic imaging, we analyzed the body temperature of *Cx. quinquefasciatus* while blood feeding. Infrared thermography has allowed us to identify a cooling strategy, evaporative cooling via the production of fluid droplets, and an overall low body temperature in comparison to the blood temperature during feeding. Understanding *Cx. quinquefasciatus’* adaptations and various strategies that they employ to reduce their body temperature while blood-feeding constitutes the first step towards the discovery of potential targets of opportunity for their control.

**Highlights:**

- Mosquitoes have evolved to cope with heat stress associated with warm blood ingestion
- *Culex quinquefasciatus* displays heterothermy while blood-feeding
- The abdominal temperature decreases due to evaporative cooling using urine droplets
- Overall, the mosquito body temperature is much cooler than the ingested blood

## 1. INTRODUCTION

Mosquitoes are the deadliest animal on the planet, killing an estimated 1 million people per year (WHO, 2020). Mosquitoes vector several pathogens responsible for devastating diseases, such as malaria, spread by *Anopheles* spp., and dengue, chikungunya, and Zika, spread primarily by *Aedes* spp (Reviewed by Reinhold *et al*., 2018; WHO, 2020). West Nile virus (WNV), which has had an increase in cases by 25% in the last few years, and other diseases causing encephalitis are mainly spread by *Culex* spp. (CDC, 2019; WHO, 2020). Vaccines and treatments have either not been developed or are limited in efficacy for these diseases, thus vector control remains the primary method of prevention (WHO 2020). *Culex quinquefasciatus* (Say, 1823) (previously *Culex pipiens fatigans*), or the southern house mosquito, is part of the *Culex pipiens* complex and inhabits tropical and subtropical regions of the World including the Americas, Africa, the Middle East, Asia and Australia (Samy *et al*., 2016). As a part of this complex, it is one of the primary vectors for WNV (Kent *et al*., 2010; Molaei *et al*., 2007), St. Louis encephalitis (SLE) (Diaz *et al*., 2013; Meyer *et al*., 1983), Western equine encephalitis (WEE) in China (Wang *et al*., 2012) and Japanese encephalitis (dos Santos Malafronte, 2003; Nitatpattana *et al*., 2005). This species also vectors an insect-specific virus, Culex flavivirus (CxFV) (Hoyos-Lopez *et al*., 2016, Farfan-Ale *et al*., 2010), which may increase the likelihood of infection and the transmission of WNV (Kent *et al*., 2010; Newman *et al*., 2011). Moreover, *Cx. quinquefasciatus* is the primary vector for the parasite *Wuchereria bancrofti*, the causative agent of lymphatic filariasis (Reviewed by Farajollahi *et al*., 2011; Triteeraprapab *et al*., 2000; Vythilingam et al, 2005). While this species is an important disease vector, it is often ignored in favor of research for *Cx. pipiens*. This could be largely due to the geographic distribution of the two species, *Cx. pipiens* occupying more northern, and, often, more wealthy countries than *Cx. quinquefasciatus* (Vinogradova, 2000). Climate change could potentially affect the distribution of *Cx. quinquefasciatus*, allowing it to spread further north into the US (Samy *et al*., 2016). This species feeds primarily on birds, taking a blood meal from mammals (typically humans and dogs) occasionally (Garcia-Rejon *et al*., 2010, Molaei *et al*., 2007). However, depending on the region the species is in and host availability, it has also been described as anthropophilic (Dixit *et al*., 2001; Hamon, 1963; Subra, 1970). This species is nocturnal and is commonly found in urban areas, feeding both endo- and exophagously (Hamon, 1963; Subra, 1970), and they will often take more than one blood meal, increasing chances for pathogen transmission (Subra, 1981). Although this is a species that feeds exclusively on warm-blooded vertebrates (particularly birds, which tend to have warmer body temperatures than humans), the mechanism by which this mosquito tolerates a warm blood meal several times in its lifetime is unknown.

Temperature is a critical abiotic factor affecting mosquito biological, behavioral and physiological processes (Reinhold *et al*., 2018). As poïkilotherms, these organisms do not maintain a stable body temperature and are dependent on the environmental temperature. All insects have an optimum temperature at which they can perform properly (Mellanby, 1939; Upshur et al., 2019). However, since environmental temperature is not consistent, animals must adjust to prevent freezing or desiccation. Insects are particularly vulnerable to temperature changes as their open circulatory and respiratory systems leave them susceptible to desiccation (Headlee, 1914; Rolandi *et al*., 2014). Many insects have evolved strategies for surviving through or avoiding cold temperatures, which range from moving to a warmer area (*i.e*., basking) (Kent *et al*., 2004) to physiological changes (*e.g*., diapause, cryoprotectants, etc.) (Heinrich, 1995) to actively moving muscles, particularly flight muscles, in order to produce heat (*i.e*., shivering) (Esch, 1988; Heinrich and Kammer, 1973). On the other hand, an ambient temperature that is too high can cause the insect to slow activity to avoid overheating and desiccation (Heinrich and Esch, 1994; Upshur *et al*., 2019). Bumblebees (*Bombus vosnesenskii*) are known to transfer heat from the thorax to the abdomen (or vice versa) in order to cool the thorax (or abdomen) when reaching a high temperature (Heinrich 1976). Additionally, Heinrich showed that honeybees (*Apis mellifera*) use evaporative cooling by regurgitating and releasing droplets through the ventral portion of the head to evaporate (1980a) and that these bees cool their thorax by transferring heat to the head, which can then cool with evaporative cooling (1980b). However, some insects, such as blood-sucking insects, have to face a sudden increase of their body temperature associated with blood intake from a warm-blooded host (Benoit et al., 2011; Beyenbach and Piermarini 2011).

Hematophagous insects have developed a necessity to feed on blood. These insects may need a blood meal because it is a sole nutrient source, like in kissing bugs (Rabinovich *et al*., 2011), or for advancement to the next life stage, like in bed bugs (Reinhardt and Siva-Jothy, 2007), or for nutrients in egg development, like in female mosquitoes (Reviewed by Adams, 1999; Gulia-Nuss et al., 2015). While blood-feeding is already a risky behavior due to host defenses (Vinauger *et al*., 2018), insects that feed on warm blooded vertebrates must face the risk of thermal stress in addition to evading host defenses (Beyenbach and Piermarini, 2011). However, some species have developed various coping mechanisms to avoid overheating during the sudden intake of a hot liquid (Benoit et al., 2019). Insects can cool their body temperature during feeding using countercurrent heat exchange, like in *Rhodnius prolixus (*Lahondère *et al*., 2017), or evaporative cooling, as in *Anopheles stephensi* (Lahondère and Lazzari, 2012). Another method consists of synthesizing heat shock proteins (HSPs) to prevent damage caused by heat stress post-feeding, which is used in *Aedes aegypti* (Benoit *et al*., 2011; Lahondère and Lazzari, 2012), and *Cx. pipiens* (Benoit *et al*., 2011). However, to our knowledge, nothing is known about the response to blood intake in the closely related species, *Cx. quinquefasciatus*. In this study, we seek to determine how *Cx. quinquefasciatus* copes with the thermal stress associated with the intake of a warm blood meal using experimental blood feeding coupled with thermography. This allowed us to follow the evolution of the body temperature of the females while feeding and identify possible cooling mechanism.

## 2. MATERIALS AND METHODS

### 2.1. Mosquitoes

*Cx. quinquefasciatus* Say (Diptera: Culicidae) eggs (NR-43025, *Culex quinquefasciatus*, strain JHB) were received from the Center for Disease Control and Prevention (Atlanta, GA). Mosquitoes were reared from eggs, which were collected as rafts from the previous generation and hatched in a larval tray (BioQuip Products, Rancho Dominguez, CA) containing deionized water and Hikari First Bites powdered fish food (Kyori Food Industries, Japan). Mosquitoes were maintained in a climatic chamber at 26 ± 1°C, 70% humidity and a 12:12 light cycle. The larvae were maintained in the trays and were isolated within 24 hours of pupation and then transferred to containers (BioQuip Products, Rancho Dominguez, CA) until emergence. The mosquitoes had *ad libitum* access to cotton balls soaked in 10% sucrose solution, which was removed before the experiments to increase the females’ motivation to take a blood-meal. The females used for the study were 6-10 days old and starved for 24 hours before testing.

### 2.2. Experimental Blood Feeding

Mosquitoes were released into a covered 8 × 8 × 8” metal collapsible cage (BioQuip Products, Rancho Dominguez, CA) and allowed to feed on adult bovine blood (Lampire Biological Laboratories, Pipersville, PA). Blood was placed into a water bath-heated glass blood feeder (D.E. Lillie Glassblowers, Atlanta, GA, USA), covered with a Parafilm membrane (Bemis Company, Inc, Neenah, WI, USA). The water bath (Jublo Corio Open Heating Bath Circulators, Thomas Scientific, Swedesboro, NJ) was heated to 41°C, and the blood was warmed to 37 ∓ 1°C before mosquitoes were released in the cage (Fig. 1). The mosquitoes that fed on the membrane were filmed using a FLIR T540 thermographic camera (FLIR Systems, Wilsonville, OR) at a 30 frames / second rate.

**Figure 1:**
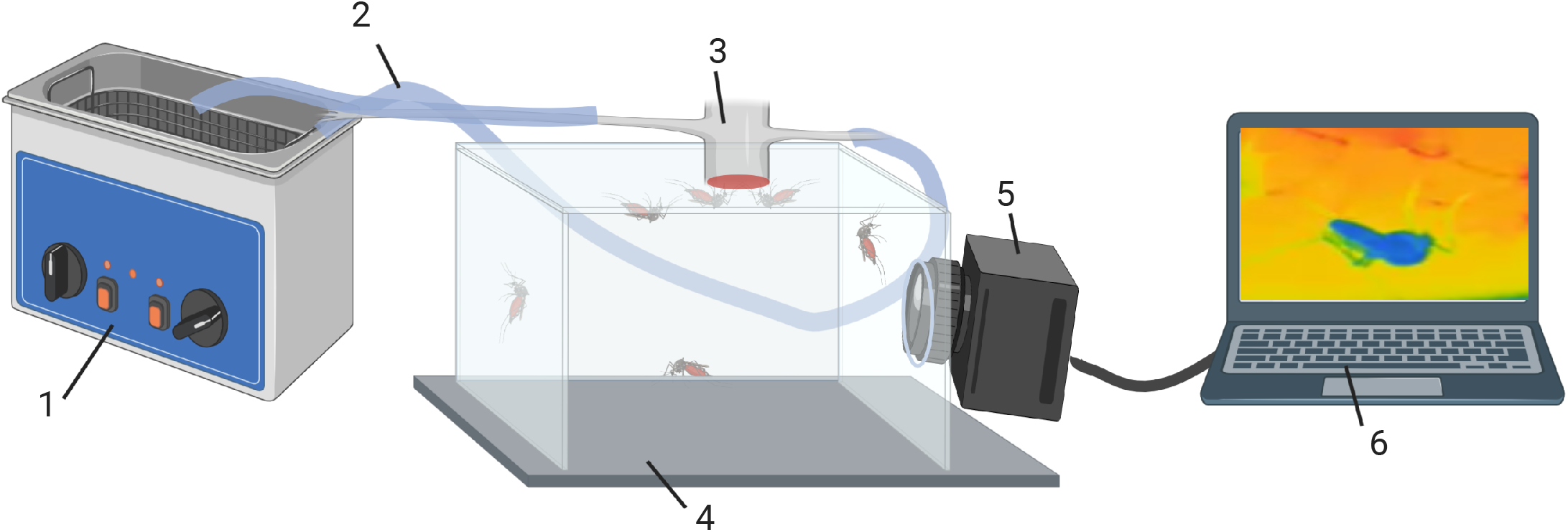
Schematic of the experimental blood feeding setup: 1. Water bath; 2. Tubing connecting water bath to blood feeder; 3. Blood feeder, containing cow’s blood; 4. Cage containing several female *Cx. quinquefasciatus* mosquitoes; 5. FLIR thermographic camera; 6. Laptop containing ResearchIR software for video analysis.

### 2.3. Video and Data Analyses

The videos (in .csq format) recorded with the thermographic camera were analyzed using the ResearchIR software (FLIR). For each mosquito, a region of interest (ROI) point was selected on the center of head, thorax, and abdomen. The software tracked the changes and evolution of the body temperature of the 3 ROIs during blood-feeding, frame by frame. For each mosquito, 10 frames were randomly selected per body segment and averaged for each mosquito. These data were then pooled (N = 10) and were compared with pairwise Student *t*-tests using the software R (R software, version 3.6.3).

## 3. RESULTS

Prior to landing on the blood feeder, the body temperature (T_*b*_) of the mosquito matched the ambient temperature (T_*a*_) (Figure 2A), and the temperature of the head (T_*h*_), the thorax (T_*th*_), and the abdomen (T_*ab*_) were all within one degree Celsius of each other (Figure 2A’). After landing, the mosquito quickly warmed up purely based on contact with the feeder. Once feeding began, T_*h*_, T_*th*_, and T_*ab*_ gradually increased (Figure 2B), and the temperatures of the three sections were increasingly different, a phenomenon known as heterothermy (Figure 2B’). As feeding continues, T_*h*_ (30.8 ±1.1 °C) was significantly higher than T_*th*_ (29.8 ±1.1 °C) (Student *t* test, t = 7.4867, df = 9, p-value = <0.001) and T_*ab*_ (28.8 ±1.1 °C) (Student *t* test, t = 10.377, df = 9, p-value = <0.001). T*th* and T_*ab*_ were also significantly different from one another (Student *t* test, t = 6.0507, df = 9, p-value = <0.001) (Figures 3 and 4). Interestingly, *Cx. quinquefasciatus’* T_*h*_ was lower than other hematophagous insects when feeding on blood at the same temperature (*i.e*., 37°C) (Table 1). In some mosquitoes, we also noticed the excretion of droplets at the end of the abdomen during feeding (*i.e*., prediuresis), some of which remained attached to the tip of the abdomen (Figure 5A). Retaining the droplet resulted in a decrease in T_*ab*_ (Figure 5B). After the mouthparts were retracted from the feeder membrane, T_*b*_ gradually decreased and began to return to T_*a*_ (Figure 2C). After feeding stopped, T_*h*_, T*th*, and T_*ab*_ were close to ambient temperature (Figure 2C’). We also measured the duration of feeding in *Cx. quinquefasciatus* (Table 1). At greater than 3 minutes on average, this species took slightly longer than most other warm-blood feeding mosquitoes to imbibe a blood meal at 37°C. However, it remains much less than the mosquito *Cx. territans*, which primarily feeds on cold-blooded animals including frogs and snakes, or other hematophagous insects (Table 1).

**Figure 2:**
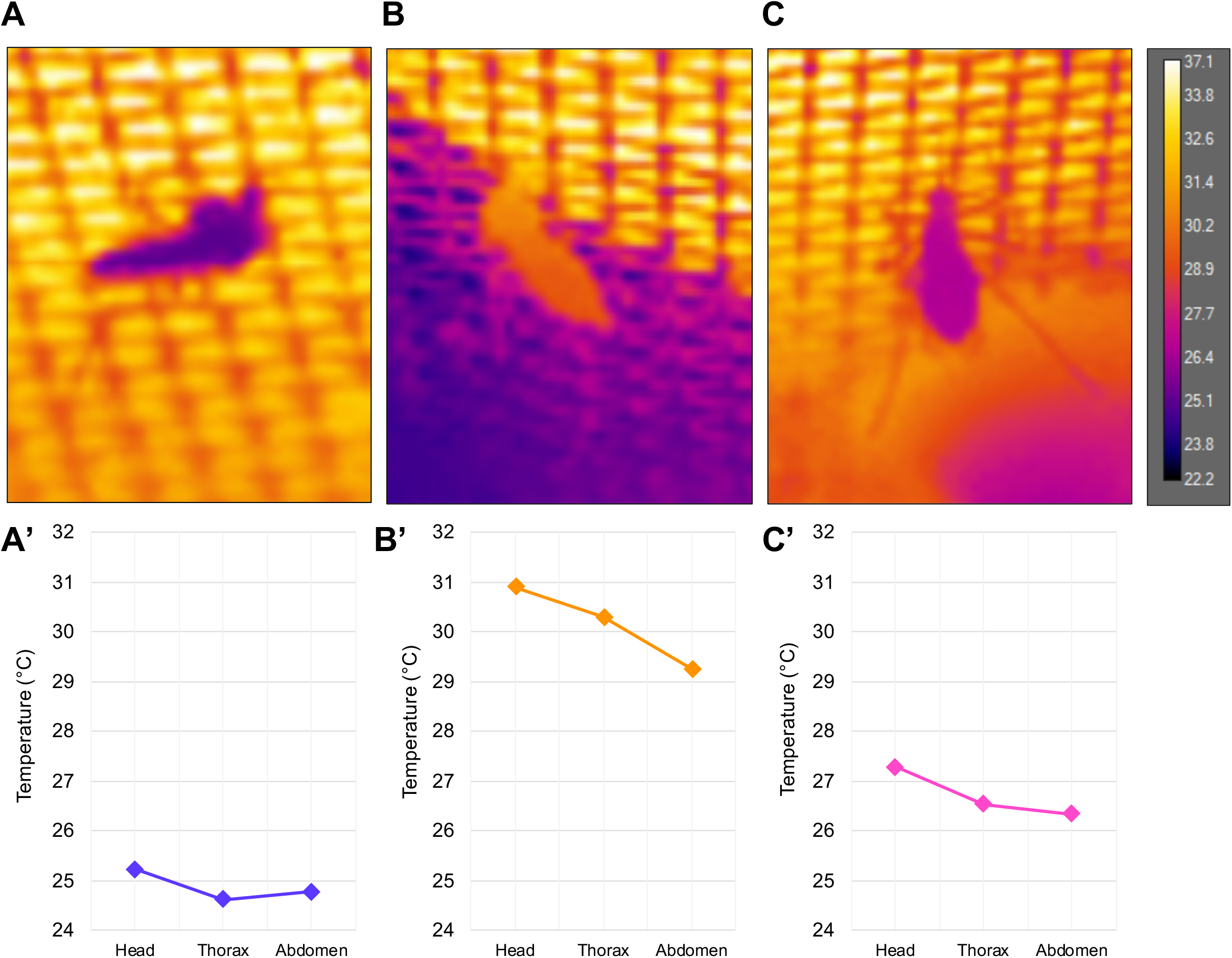
Thermographic images of *Cx. quinquefasciatus* female mosquitoes taken shortly after landing on the blood-feeder (**A**), during blood-feeding (*T_blood_* = 37 ± 1°C) (**B**), and after feeding (**C**) and their respective body temperatures (**A’**, **B’**, **C’**).

**Figure 3:**
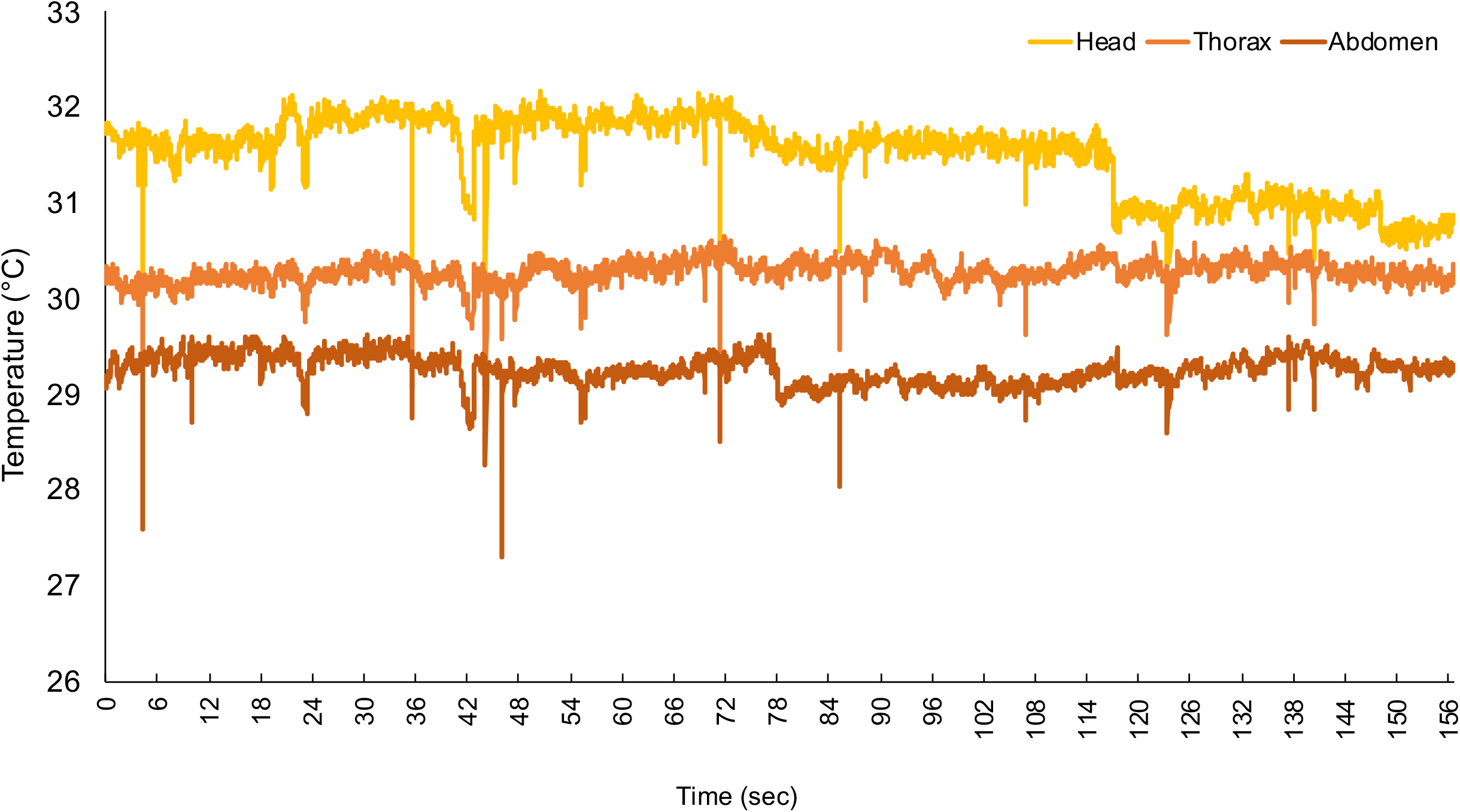
Evolution of the temperature of the head (yellow), thorax (orange) and abdomen (red) of a *Cx. quinquefasciatus* female during feeding on a blood-meal at 37 ± 1°C. Recordings were conducted at 30 frames / sec.

**Figure 4:**
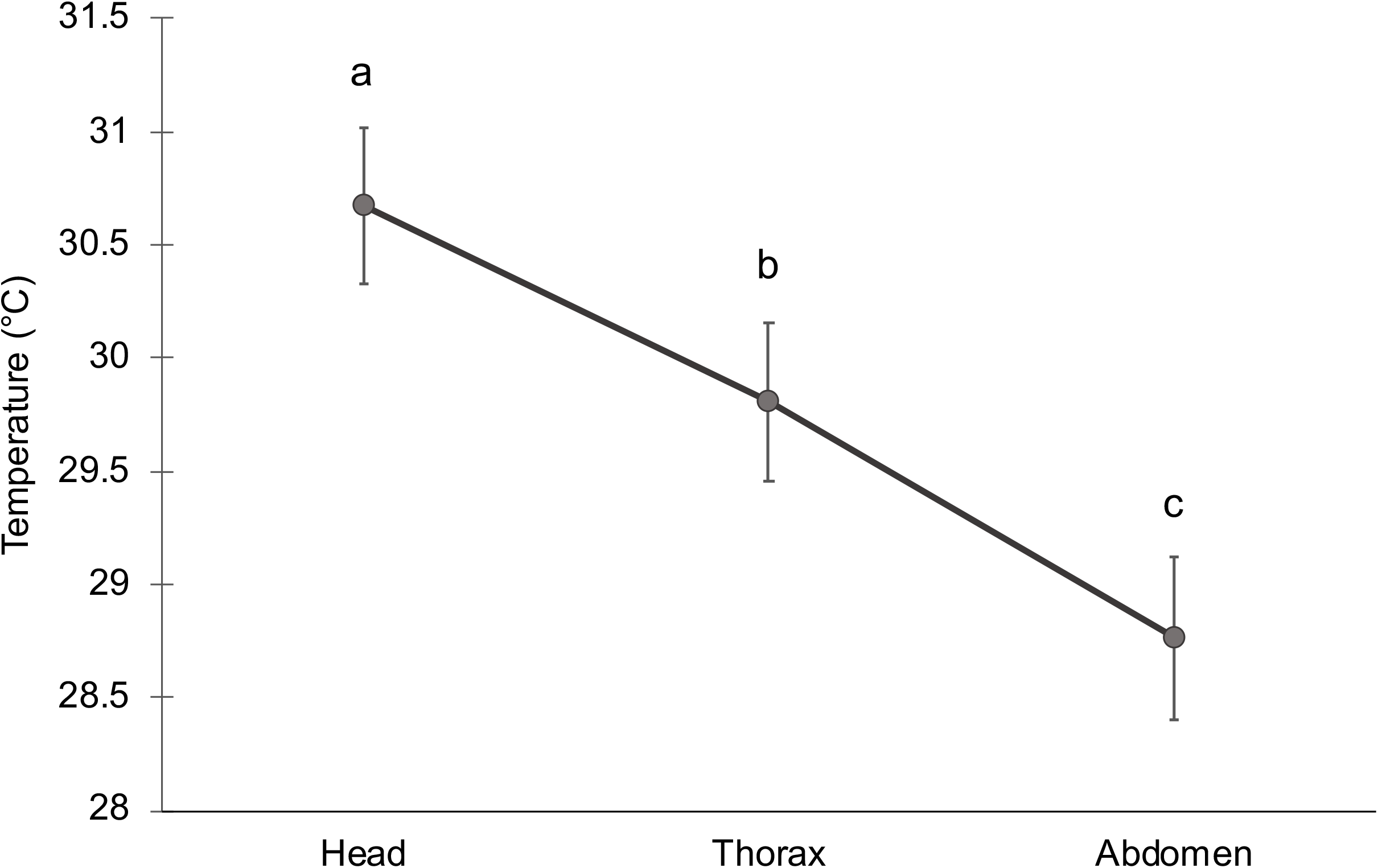
Average temperatures of the head, thorax and abdomen of *Cx quinquefasciatus* females (N = 10) feeding on a blood-meal at 37 ± 1°C. Vertical bars represent the standard error of the mean values (S.E.M.) Letters above the error bars indicate statistical differences (Student *t*-tests, p < 0.001).

**Figure 5:**
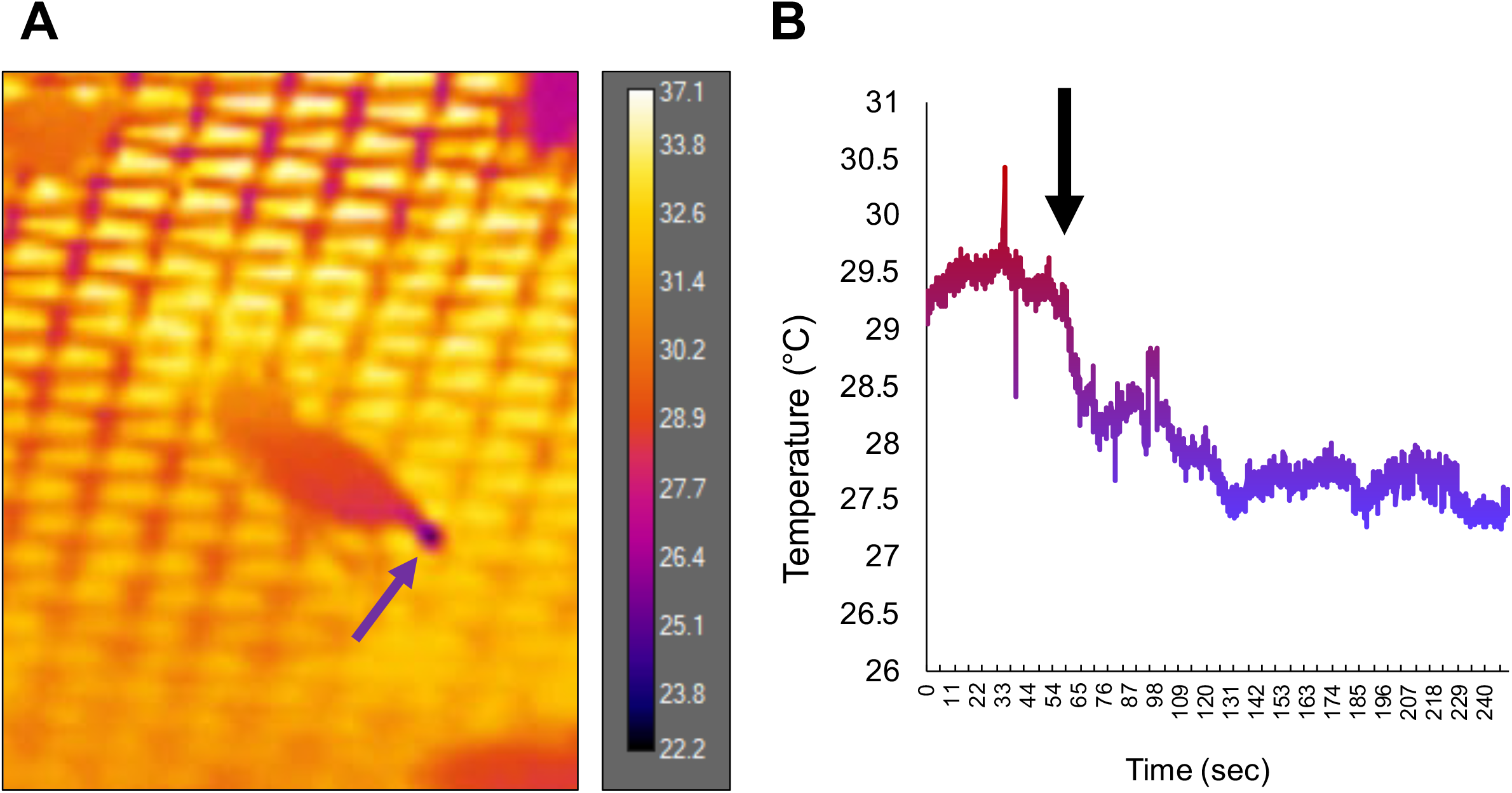
**A.** Thermographic image of a *Cx. quinquefasciatus* female during feeding on a blood-meal at 37 ± 1°C producing and maintaining a droplet of urine at the tip of its abdomen, indicated by the arrow. **B.** Evolution of the temperature of its abdominal temperature before the droplet emission / retention (red) and after (blue). The emission of the droplet occurs 60 seconds after starting feeding (black arrow) and remains attached until the female takes off.

**Table 1:**
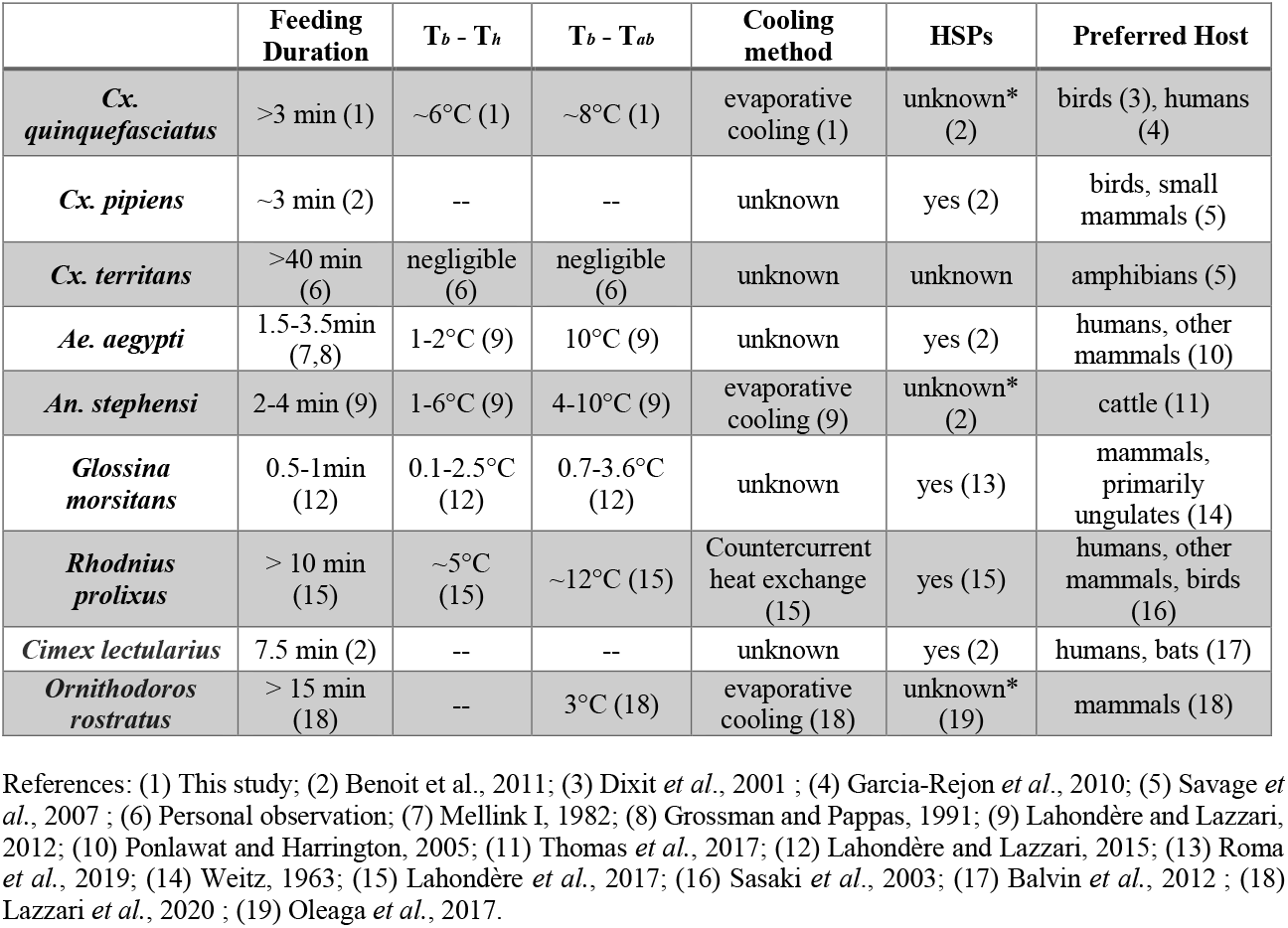
Summary of the feeding behavior and physiological characteristics of several blood-sucking arthropods. (*) indicates that HSPs are synthesized by a closely related species. Numbers in parentheses indicate the reference of the studies.

## 4. DISCUSSION

In the present study, we showed that *Cx. quinquefasciatus* displays heterothermy, a temperature gradient along the body segments, while feeding on warm blood. This has been observed in several hematophagous insect species, but the underlying mechanisms can vary (Benoit *et al*., 2019; Lahondère and Lazzari, 2013). Countercurrent heat exchange is mostly seen in the literature as a method of heat conservation (*e.g*., Casey, 1988; Heinrich, 1995), but Lahondère *et al*. (2017) showed that the kissing bug, *Rhodnius prolixus*, uses countercurrent heat exchange to cool its abdomen down while feeding. Cool hemolymph is pumped from the abdomen to the head through the heart and aorta, which comes into contact with the esophagus in the head. The hot ingested blood circulating into the esophagus transfers some of its heat to the hemolymph in the aorta, which helps to cool the blood before it reaches the crop. Another strategy, evaporative cooling, consists of excreting and retaining droplets of fluid in order to cool down (Heinrich, 1980a; Prange, 1996). Several hematophagous arthropods, including sandflies (Sadlova *et al*., 2013), mosquitoes (Lahondère and Lazzari, 2012) and ticks (Lazzari et al., 2020), use this method to cool down during blood intake. Mosquitoes use prediuresis droplets for evaporative cooling, in which the insect excretes a droplet composed of fresh blood and urine and holds it at the tip of the abdomen to cool down (Lahondère and Lazzari, 2012). In *An. stephensi*, prediuresis is observed during blood intake, and keeping the droplet to perform evaporative cooling occurs in most individuals (Lahondère and Lazzari, 2012; 2013). In the present study, we noted only a small percentage of *Cx. quinquefasciatus* displaying this behavior. Future experiments in which mosquitoes are fed with blood at different temperatures will inform whether evaporative cooling occurrence is dependent on the temperature of the host blood.

While most species of hematophagous insects have a head temperature close to the temperature of the blood while feeding, here, we noted that *Cx. quinquefasciatus’* head temperature was much lower than that of the blood. Lahondère and Lazzari showed a few degree difference between the blood temperature and the mosquito head temperature (in this case, *Ae. aegypti* and *An. stephensi*) (2012) or the tsetse fly *Glossina morsitans* (<2°C difference) (2015), whereas we observed a much larger difference (~6°C on average). This suggests that *Cx. quinquefasciatus* has developed a way to cool blood down before it reaches the head. While this has not been seen in mosquitoes, cooling of the head has been seen in honeybees, which use evaporative cooling by releasing droplets from the ventral portion of the head (Heinrich, 1980a). We put forth the hypothesis that the ingestion pumps in *Cx. quinquefasciatus* may have a specialized function that may play a role in this. Investigating the pumps’ anatomy and function during blood ingestion will provide insights into the mechanisms underlying heterothermy in *Cx. quinquefasciatus* (Kikuchi *et al*., 2018; Lahondère *et al*., 2017). Another possibility is that *Cx. quinquefasciatus* could be controlling its body temperature by intaking the blood more slowly than other mosquito species. Chadee *et al*. (2002) found both fast and slow feeders in *Ae. aegypti* (less than or greater than 2 minutes, respectively), whereas *Cx. quinquefasciatus* took an average of 3 minutes to take a full meal (Table 1). This slower feeding time could allow the blood to lose heat through the cuticle of the stylets as the mosquito ingests the blood meal.

To avoid heat stress associated with the ingestion of a warm blood meal, mosquitoes can shift their host preference and select relatively cooler blood to feed on. We made some observations of mosquitoes seemingly preferring to feed on the outer rim of the blood feeder, where the blood is slightly cooler. This behavior has been seen with hosts as well, as Oduola and Awe (2006) found that *Cx. quinquefasciatus* prefers to feed on the foot and ankle rather than the calf and thigh, which could be due to odorants produced by the foot, but it may also be affected by the lower temperature of the extremities (Aminoff *et al*., 2018). Host or biting site selection, *i.e*., preferring to feed on the lowest temperature area available, may also contribute to evaporative cooling droplet formation, where mosquitoes forced to feed on warmer blood may be more likely to form a droplet. Comparing the feeding behavior of *Cx. quinquefasciatus* at varying temperatures will allow us to test this hypothesis. It is worth mentioning that when we analyzed the feeding activity of the closely related cold-blooded feeding mosquito, *Culex territans*, we found that this species takes 13 times longer to feed to repletion than *Cx. quinquefasciatus* (Table 1).

*Cx. quinquefasciatus* may also synthesize heat shock proteins (HSPs), which typically function as chaperone proteins to prevent damage to existing proteins during times of stress, to recover from ingestion of a hot blood meal. Because the production of HSPs in response to thermal stress is a highly conserved physiological response to heat stress throughout the animal kingdom (Pereira *et al*., 2017), and *Cx. pipiens* has been shown to use these proteins after ingesting a hot blood-meal (Benoit *et al*., 2011), we suggest that *Cx. quinquefasciatus* might also respond to heat stress in this way. Tsetse flies have been shown to synthesize HSPs (Roma *et al*., 2019) and several mosquito species, including *Ae. aegypti* and *Culex pipiens*, use this method to recover from the stress caused by rapid intake of a warm meal (Benoit *et al*., 2011; Lahondère and Lazzari, 2012). In kissing bugs, it has been shown that both exposure to heat and blood feeding trigger the synthesis of HSP70 (Paim et al. 2016). Moreover, several species of hard and soft ticks also synthesize HSPs to recover from heat stress (Guilfoile and Packila, 2004; Busby et al. 2012; Oleaga et., 2017). It is thus likely that *Cx. quinquefasciatus* is synthesizing HSPs in response to the intake of a hot blood meal in addition to evaporative cooling and possibly other cooling mechanisms yet unknown.

## 5. CONCLUSIONS

*Cx. quinquefasciatus* plays a large role in disease transmission across the world. To our knowledge, this study is the first focusing on the thermal biology of this mosquito species. We showed that this mosquito cools down during blood-feeding in part via evaporative cooling of urine droplets. Understanding the biology of this mosquito, particularly of its feeding behavior and physiology, can lead to more integrative pest management methods in order to control this vector.

## Acknowledgements

We are grateful to members of the Vinauger lab and the Lahondère lab for mosquito colony care. The following reagent was provided by Centers for Disease Control and Prevention for distribution by BEI Resources, NIAID, NIH: *Culex quinquefasciatus*, Strain JHB, Eggs, NR-43025. Figure 1 was created with BioRender.com. Funding sources: The Global Change Center (C.L. and R.S), the Fralin Life Sciences Institute (C.L.) and the Department of Biochemistry (C.L.) at Virginia Tech.

## References

Adams, T. S. (1999). Hematophagy and hormone release. Annals of the Entomological Society of America, 92(1), 1–13.

Aminoff, M. J., Boller, F., Swaab, D. F., Romanovsky, A. A., Garami, A. et al. (2018). Thermoregulation: from basic neuroscience to clinical neurology. ed. Andrej A. Romanovsky, Elsevier. 496pp.

Balvín, O., Munclinger, P., Kratochvíl, L., & Vilímová, J. (2012). Mitochondrial DNA and morphology show independent evolutionary histories of bedbug *Cimex lectularius* (Heteroptera: Cimicidae) on bats and humans. Parasitology Research, 111(1), 457–469.

Benoit, J. B., Lopez-Martinez, G., Patrick, K. R., Phillips, Z. P., Krause, T. B. et al. (2011). Drinking a hot blood meal elicits a protective heat shock response in mosquitoes. Proceedings of the National Academy of Sciences, 108(19), 8026–8029.

Benoit, J. B., Lazzari, C. R., Denlinger, D. L., & Lahondère, C. (2019). Thermoprotective adaptations are critical for arthropods feeding on warm-blooded hosts. Current opinion in insect science, 34, 7–11.

Beyenbach, K. W., & Piermarini, P. M. (2011). Transcellular and paracellular pathways of transepithelial fluid secretion in Malpighian (renal) tubules of the yellow fever mosquito *Aedes aegypti*. Acta Physiologica, 202(3), 387–407.

Busby, A. T., Ayllón, N., Kocan, K. M., Blouin, E. F., De La Fuente, G. et al. (2012). Expression of heat shock proteins and subolesin affects stress responses, *Anaplasma phagocytophilum* infection and questing behaviour in the tick, *Ixodes scapularis*. Medical and Veterinary Entomology, 26(1), 92–102.

Casey, T. M. (1988). Thermoregulation and heat exchange. Advances in Insect Physiology, 20, 119–146.

Center for Disease Control and Prevention, 2019. West Nile virus and other domestic nationally notifiable arboviral diseases--United States, 2018. https://www.cdc.gov/mmwr/volumes/68/wr/mm6831a1.htm?s_cid=mm6831a1_e&deliveryName=USCDC_921-DM6573.

Chadee, D. D., Beier, J. C., & Mohammed, R. T. (2002). Fast and slow blood-feeding durations of *Aedes aegypti* mosquitoes in Trinidad. Journal of Vector Ecology, 27, 172–177.

Diaz, L. A., Flores, F. S., Beranek, M., Rivarola, M. E., Almirón, W. R. et al. (2013). Transmission of endemic St Louis encephalitis virus strains by local *Culex quinquefasciatus* populations in Cordoba, Argentina. Transactions of the Royal Society of Tropical Medicine and Hygiene, 107(5), 332–334.

Dixit, V., Gupta, A. K., Kataria, O. M., & Prasad, G. B. (2001). Host preference of *Culex quinquefasciatus* in Raipur city of Chattisgarh state. The Journal of Communicable Diseases, 33(1), 17–22.

dos Santos Malafronte, R., Calvo, E., James, A. A., & Marinotti, O. (2003). The major salivary gland antigens of *Culex quinquefasciatus* are D7-related proteins. Insect biochemistry and molecular biology, 33(1), 63–71.

Esch, H. (1988). The effects of temperature on flight muscle potentials in honeybees and cuculiinid winter moths. Journal of Experimental Biology, 135(1), 109–117.

Farfan-Ale, J. A., Lorono-Pino, M. A., Garcia-Rejon, J. E., Soto, V., Lin, M. et al. (2010). Detection of flaviviruses and orthobunyaviruses in mosquitoes in the Yucatan Peninsula of Mexico in 2008. Vector-Borne and Zoonotic Diseases, 10(8), 777–783.

Farajollahi, A., Fonseca, D. M., Kramer, L. D., & Kilpatrick, A. M. (2011). “Bird biting” mosquitoes and human disease: a review of the role of *Culex pipiens* complex mosquitoes in epidemiology. Infection, genetics and evolution, 11(7), 1577–1585.

Garcia-Rejon, J. E., Blitvich, B. J., Farfan-Ale, J. A., Loroño-Pino, M. A., Chim, W. A. C. et al. (2010). Host-feeding preference of the mosquito, *Culex quinquefasciatus*, in Yucatan State, Mexico. Journal of Insect Science, 10.

Grossman, G. L., & Pappas, L. G. (1991). Human skin temperature and mosquito (Diptera: Culicidae) blood feeding rate. Journal of medical entomology, 28(3), 456–460.

Guilfoile, P. G., & Packila, M. (2004). Identification of four genes expressed by feeding, female *Ixodes scapularis* including three with sequence similarity to previously recognized genes. Experimental & applied acarology, 52(1-2), 103–110.

Gulia-Nuss, M., Elliot, A., Brown, M. R., & Strand, M. R. (2015). Multiple factors contribute to anautogenous reproduction by the mosquito *Aedes aegypti*. Journal of insect physiology, 82, 8–16.

Hamon, J. (1963). Les moustiques anthropophiles de la région de Bobo Dioulasso, République de Haute Volta. Annals de la Société Entomologique de France. 85–145.

Headlee, T. J. (1914). Some data on the effect of temperature and moisture on the rate of insect metabolism. Journal of Economic Entomology, 7, 413.

Heinrich, B. (1976). Heat exchange in relation to blood flow between thorax and abdomen in bumblebees. Journal of Experimental Biology, 64(3), 561–585.

Heinrich, B. (1980a). Mechanisms of body-temperature regulation in honeybees, *Apis mellifera:* I. Regulation of head temperature. Journal of Experimental Biology, 85(1), 61–72.

Heinrich, B. (1980b). Mechanisms of body-temperature regulation in honeybees, *Apis mellifera*. II. Regulation of thoracic temperature at high air temperatures. Journal of Experimental Biology, 85, 73–87.

Heinrich, B. (1995). Insect thermoregulation. Endeavour, 19(1), 28–33.

Heinrich, B., & Esch, H. (1994). Thermoregulation in bees. American Scientist, 82(2), 164–170.

Heinrich, B., & Kammer, A. E. (1973). Activation of the fibrillar muscles in the bumblebee during warm-up, stabilization of thoracic temperature and flight. Journal of Experimental Biology, 58(3), 677–688.

Hoyos-López, R., Suaza-Vasco, J., Rúa-Uribe, G., Uribe, S., & Gallego-Gómez, J. C. (2016). Molecular detection of flaviviruses and alphaviruses in mosquitoes (Diptera: Culicidae) from coastal ecosystems in the Colombian Caribbean. Memórias do Instituto Oswaldo Cruz, 111(10), 625–634.

Kemp, D. J., & Krockenberger, A. K. (2004). Behavioural thermoregulation in butterflies: the interacting effects of body size and basking posture in *Hypolimnas bolina* (L.)(Lepidoptera: Nymphalidae). Australian journal of Zoology, 52(3), 229–239.

Kent, R. J., Crabtree, M. B., & Miller, B. R. (2010). Transmission of West Nile virus by *Culex quinquefasciatus* (Say) infected with Culex Flavivirus Izabal. PLoS neglected tropical diseases, 4(5).

Kikuchi, K., Stremler, M. A., Chatterjee, S., Lee, W. K., Mochizuki, O. et al. (2018). Burst mode pumping: A new mechanism of drinking in mosquitoes. Scientific reports, 8(1), 1–15.

Lahondère, C., & Lazzari, C. R. (2012). Mosquitoes cool down during blood feeding to avoid overheating. Current biology, 22(1), 40–45.

Lahondère, C., & Lazzari, C. R. (2013). Thermal stress and thermoregulation during feeding in mosquitoes. In Anopheles mosquitoes – New insights into malaria vectors. IntechOpen.

Lahondère, C., & Lazzari, C. R. (2015). Thermal effect of blood feeding in the telmophagous fly *Glossina morsitans morsitans*. Journal of thermal biology, 48, 45–50.

Lahondère, C., Insausti, T. C., Paim, R. M., Luan, X., Belev, G. et al. (2017). Countercurrent heat exchange and thermoregulation during blood-feeding in kissing bugs. Elife, 6, e26107.

Lazzari, C.R., Fauquet, A., Lahondère, C, Araujo, R. & M.H. Pereira. Soft ticks perform evaporative cooling during blood-feeding (2020). BioRxiv. https://doi.org/10.1101/2020.06.30.180968.

Mellanby, K. (1939). Low temperature and insect activity. Proceedings of the Royal Society of London. Series B-Biological Sciences, 127(849), 473–487.

Mellink, J. J., Poppe, D. M. C., & Van Duin, G. J. T. (1982). Factors affecting the blood-feeding process of a laboratory strain of *Aedes aegypti* on rodents. Entomologia Experimentalis et Applicata, 31(2-3), 229–238.

Meyer, R. P., Hardy, J. L., & Presser, S. B. (1983). Comparative vector competence of *Culex tarsalis* and *Culex quinquefasciatus* from the Coachella, Imperial, and San Joaquin Valleys of California for St. Louis encephalitis virus. The American journal of tropical medicine and hygiene, 32(2), 305–311.

Molaei, G., Andreadis, T. G., Armstrong, P. M., Bueno Jr, R., Dennett, J. A. et al. (2007). Host feeding pattern of *Culex quinquefasciatus* (Diptera: Culicidae) and its role in transmission of West Nile virus in Harris County, Texas. The American journal of tropical medicine and hygiene, 77(1), 73–81.

Newman, C. M., Cerutti, F., Anderson, T. K., Hamer, G. L., Walker, E. D. et al. (2011). Culex flavivirus and West Nile virus mosquito coinfection and positive ecological association in Chicago, United States. Vector-Borne and Zoonotic Diseases, 11(8), 1099–1105.

Nitatpattana, N., Apiwathnasorn, C., Barbazan, P., Leemingsawat, S., Yoksan, S. et al. (2005). First isolation of Japanese encephalitis from *Culex quinquefasciatus* in Thailand. Southeast Asian journal of tropical medicine and public health, 36(4), 875.

Oduola, A. O., & Awe, O. O. (2006). Behavioural biting preference of *Culex quinquefasciatus* in human host in Lagos metropolis Nigeria. Journal of vector borne diseases, 43(1), 16.

Oleaga, A., Obolo-Mvoulouga, P., Manzano-Román, R., & Pérez-Sánchez, R. (2017). Functional annotation and analysis of the *Ornithodoros moubata* midgut genes differentially expressed after blood feeding. Ticks and tick-borne diseases, 8(5), 693–708.

Paim, R. M., Araujo, R. N., Leis, M., Sant’Anna, M. R., Gontijo, N. F. et al. (2016). Functional evaluation of Heat Shock Proteins 70 (HSP70/HSC70) on *Rhodnius prolixus* (Hemiptera, Reduviidae) physiological responses associated with feeding and starvation. Insect Biochemistry and Molecular Biology, 77, 10–20.

Pereira, M. H., Paim, R. M., Lahondère, C., & Lazzari, C. R. (2017). Heat Shock Proteins and Blood-Feeding in Arthropods. In Heat Shock Proteins in Veterinary Medicine and Sciences (pp. 349–359). Springer, Cham.

Ponlawat, A., & Harrington, L. C. (2005). Blood feeding patterns of *Aedes aegypti* and *Aedes albopictus* in Thailand. Journal of medical entomology, 42(5), 844–849.

Prange, H. D. (1996). Evaporative cooling in insects. Journal of Insect Physiology, 42(5), 493–499.

R: A language and environment for statistical computing. R Foundation for Statistical Computing, Vienna, Austria. URL http://www.R-project.org/.

Rabinovich, J. E., Kitron, U. D., Obed, Y., Yoshioka, M., Gottdenker, N. et al. (2011). Ecological patterns of blood-feeding by kissing-bugs (Hemiptera: Reduviidae: Triatominae). Memorias Do Instituto Oswaldo Cruz, 106(4), 479–494.

Reinhardt, K., & Siva-Jothy, M. T. (2007). Biology of the bed bugs (Cimicidae). Annual Review of Entomology, 52, 351–374.

Reinhold, J. M., Lazzari, C. R., & Lahondère, C. (2018). Effects of the environmental temperature on *Aedes aegypti* and *Aedes albopictus* mosquitoes: a review. Insects, 9(4), 158.

Rolandi, C., Iglesias, M. S., & Schilman, P. E. (2014). Metabolism and water loss rate of the haematophagous insect *Rhodnius prolixus:* effect of starvation and temperature. Journal of Experimental Biology, 217(24), 4414–4422.

Roma, J. S., D’souza, S., Somers, P. J., Cabo, L. F., Farsin, R., Aksoy, S., … & Weiss, B. L. (2019). Thermal stress responses of *Sodalis glossinidius*, an indigenous bacterial symbiont of hematophagous tsetse flies. PLoS neglected tropical diseases, 13(11), e0007464.

Sadlova, J., Reishig, J., & Volf, P. (2013). Prediuresis in female *Phlebotomus* sandflies (Diptera: Psychodidae). European Journal of Entomology, 95(4), 643–647.

Samy, A. M., Elaagip, A. H., Kenawy, M. A., Ayres, C. F., Peterson, A. T. et al. (2016). Climate change influences on the global potential distribution of the mosquito *Culex quinquefasciatus*, vector of West Nile virus and lymphatic filariasis. PloS one, 11(10).

Sasaki, H., Rosales, R., & Tabaru, Y. (2003). Host feeding profiles of *Rhodnius prolixus* and *Triatoma dimidiata* in Guatemala (Hemiptera: Reduviidae: Triatominae). Medical Entomology and Zoology, 54(3), 283–289.

Savage, H. M., Aggarwal, D., Apperson, C. S., Katholi, C. R., Gordon, E. et al., (2007). Host choice and West Nile virus infection rates in blood-fed mosquitoes, including members of the *Culex pipiens* complex, from Memphis and Shelby County, Tennessee, 2002–2003. Vector-Borne and Zoonotic Diseases, 7(3), 365–386.

Subra, R. (1970) Etudes écologiques sur *Culex pipiens fatigans* Wiedemann, 1828, (Diptera, Culicidae) dans une zone urbaine de savane soudanienne ouest-africaine. Lieux de repos des adultes. Cahiers ORSTOM. Série Entomologie Médicale et Parasitologie. 8, 353–376.

Subra, R. (1981). Biology and control of *Culex pipiens quinquefasciatus*\* Say, 1823 (Diptera, Culicidae) with special reference to Africa. International Journal of Tropical Insect Science, 1(4), 319–338.

Thomas, S., Ravishankaran, S., Justin, N. J. A., Asokan, A., Mathai, M. T. et al. (2017). Resting and feeding preferences of *Anopheles stephensi* in an urban setting, perennial for malaria. Malaria journal, 16(1), 1–7.

Triteeraprapab, S., Kanjanopas, K., Suwannadabba, S., Sangprakarn, S., Poovorawan, Y., et al. (2000). Transmission of the nocturnal periodic strain of *Wuchereria bancrofti* by *Culex quinquefasciatus*: establishing the potential for urban filariasis in Thailand. Epidemiology & Infection, 125(1), 207–212.

Upshur, I. F., Bose, E. A., Hart, C., & Lahondère, C. (2019). Temperature and Sugar Feeding Effects on the Activity of a Laboratory Strain of *Aedes aegypti*. Insects, 10(10), 347.

Vinauger, C., Lahondère, C., Wolff, G. H., Locke, L. T., Liaw, J. E. et al. (2018). Modulation of host learning in *Aedes aegypti* mosquitoes. Current Biology, 28(3), 333–344.

Vinogradova, E. B. (2000). *Culex pipiens pipiens* mosquitoes: taxonomy, distribution, ecology, physiology, genetics, applied importance and control (No. 2). Pensoft Publishers.

Vythilingam, I., Huat, T. C., & Ahmad, N. W. (2005). Research Note Transmission potential of *Wuchereria bancrofti* by *Culex quinquefasciatus* in urban areas of Malaysia. Tropical biomedicine, 22(1), 83–85.

Wang, Z., Zhang, X., Li, C., Zhang, Y., Xing, D. et al. (2012). Vector competence of five common mosquito species in the People’s Republic of China for Western equine encephalitis virus. Vector-Borne and Zoonotic Diseases, 12(7), 605–608.

Weitz, B. (1963). The feeding habits of *Glossina*. Bulletin of the World Health Organization, 28(5-6), 711.

World Health Organization (2020). Mosquito-borne diseases. https://www.who.int/neglected_diseases/vector_ecology/mosquito-borne-diseases/en/.

